# Evaluation of Tissue-Engineered Blood Vessels as Three-Dimensional In Vitro Testing System in Cardiovascular Research and Device Approval

**DOI:** 10.1101/2025.08.26.672497

**Authors:** Diana M. Rojas-González, Frederic Wolf, Nicole Schaaps, Roberta A. Florescu, Carolina Bienzeisler, Rahma Shahin, Pakhwan Nilcham, Felix Vogt, Stefan Jockenhoevel, Anne Turoni-Glitz, Petra Mela

## Abstract

**Background:** Disturbed crosstalk between endothelial cells (ECs) and vascular smooth muscle cells (SMCs) has an important role in atherosclerosis and restenosis after vascular intervention, however, the exact pathomechanisms are incompletely understood. Current preclinical testing models do not adequately recapitulate the complexity of human arteries. Here, we present tissue-engineered blood vessels (TEBVs) as a novel *in vitro* model and validate it for intimal hyperplasia.

**Methods:** TEBVs fabricated from SMC suspended in fibrin gel, supported by a textile mesh, were seeded with ECs at various concentrations and subjected to arterial flow conditions in a bioreactor. In addition, TEBVs underwent plain old balloon angioplasty (POBA) and implantation of bare metal stents (BMS) and drug-eluting stents (DES) at day 7 after fabrication. TEBVs were dynamically conditioned in a bioreactor for 21 days in total and monitored by optical coherence tomography.

**Results:** TEBVs with absent or incomplete endothelial layer exhibited thicker vessel walls, more disorganized and misaligned collagen, and increased cellular proliferation compared with completely endothelialized TEBVs. POBA and stent implantation were feasible 7 days after TEBV fabrication. At 14 days post-intervention, POBA-treated TEBVs exhibited significantly thicker vessel walls than untreated controls and stented TEBVs, whereas stented TEBVs showed greater lumen diameters than unstented TEBVs. Endothelial strut coverage was significantly higher in BMS-treated compared with DES-treated TEBVs. Over the course of the conditioning period, levels of IL-6, IL-8, and MCP-1 were highest in medium samples from BMS-treated TEBVs compared to DES-treated TEBVs and compared to untreated controls.

**Conclusions:** TEBVs are a promising approach towards an *in vitro* system for the study of intimal hyperplasia. Due to their similarity in size and wall thickness to human coronary arteries, TEBVs may also serve as a platform for testing new stent designs.

**Graphical Abstract:** Tissue-engineered blood vessels (TEBV) fabricated from smooth muscle cell /fibroblast mixtures suspended in fibrin gel, supported by a textile mesh, were seeded with endothelial cells and conditioned in a bioreactor system for 21 days. Different endothelialization strategies resulted in differences in wall thickness. In addition, TEBVs underwent plain old balloon angioplasty (POBA) and stent implantation. POBA-treated TEBVs exhibited thicker vessel walls compared with non-treated TEBV controls and compared with stented TEBVs. We also observed significantly higher stent strut coverage with endothelial cells after implantation of bare metal stents (BMS) compared to drug-eluting stents (DES). TEBVs are a promising approach towards an *in-vitro* system for the study of intimal hyperplasia. Due to their similarity in size and wall thickness to human coronary arteries, TEBVs may also serve as a platform for testing new stent designs.

## Introduction

Arterial vessel walls are continuously exposed to mechanical, hemodynamic, and neurohumoral stimuli. Orchestrated by the interplay between cells within the arterial wall, both physiologic and pathologic stimuli elicit adaptive changes in structure and functionality of the vascular wall.

The integrity of the innermost layer of the vessel wall, the vascular endothelium, is a prerequisite for vascular homeostasis. Vascular endothelial cells (EC) not only act as a barrier between vessel wall and blood stream but also regulate the passage of macromolecules and control the vascular tone. The endothelium coordinates inflammation by mediating the recruitment of inflammatory cells and platelets and keeps the balance between coagulation and fibrinolysis. Disturbances of normal endothelial functions, however, may pave the way for the initiation and progression of intimal hyperplasia (IH) and atherosclerosis (1, 2).

Vascular smooth muscle cells (SMCs), the main cellular component of the vascular medial layer, exhibit remarkable plasticity. While SMCs have a contractile phenotype under physiological conditions, they can differentiate to a synthetic state, along with increased abilities to migrate and to proliferate in response to injury (3). These phenotypic and functional transitions in vascular SMCs have a major role in arterial remodeling (3).

The crosstalk between ECs and SMCs is crucial for vascular homeostasis and has critical impact in vascular disease (4–6). ECs and SMCs interact both directly by cell-to-cell contacts and indirectly via extracellular matrix (ECM) proteins or through secreted molecules and extracellular vesicles (4, 7). Importantly, a continuous release of vasoactive compounds by ECs is required to maintain SMCs in a contractile and quiescent state (5, 8–10). However, endothelial dysfunction or even loss, resulting from mechanical or chemical injury, a key event in atherogenesis (1), implies aberrant EC-SMC interactions. Furthermore, catheter-based interventional procedures such as plain old balloon angioplasty (POBA) and stent implantation inevitably lead to endothelial denudation (11–13). The consequential loss of a continuous and functional endothelial layer may result in SMC proliferation and their transition into a synthetic phenotype with subsequent accumulation of ECM proteins, mainly collagen-I and collagen III, within the vascular wall (6, 14, 15). Therefore, disturbed crosstalk between ECs and SMCs is an important etiologic factor in IH, and in the development of restenosis after POBA and stent implantation (16).

However, the exact pathomechanisms of how disturbed EC-SMC communication may trigger atherogenesis and restenosis are incompletely understood. While abundant studies have investigated the functions of ECs and SMCs as independent entities, a more accurate *in vitro* analysis of the pathophysiology of blood vessels requires co-culture systems of ECs and SMCs. A variety of co-culture setups have been introduced over time (8–10, 17–22).

Two-dimensional bilayer models of ECs and SMCs in co-culture are relatively cheap and allow for very basic investigation of EC-SMC interactions. While bilayer models are important for initial drug- and permeability testing, they are not ideal to gain full insight into cellular phenotypes and behavior under physiologic and pathologic flow and culture conditions (23–27). Preclinical animal models on the other hand are still a prerequisite for the regulatory approval of new devices and pharmaceutics. However, according to the principle of the 3R’s (replacement, reduction, and refinement), the use of animals for medical research gets more and more restricted (26). Moreover, animal research is associated with high costs. An improved *in vitro* testing environment may help reduce the number of animals used in research. Moreover, a three-dimensional tubular construct of human coronary artery-like dimensions, composed of ECs and SMCs in co-culture and amenable to vascular intervention, could enable the investigation of EC-SMC-interactions following POBA and stent implantation, thereby providing a valuable platform for preclinical testing of coronary devices such as coronary stents.

Here, we present tissue-engineered blood vessels (TEBV) fabricated from SMC/ fibroblast (FB) mixtures suspended in a matrix of fibrin gel and supported by a textile polyvinylidene difluoride (PVDF) mesh. TEBVs were seeded with ECs at different concentrations to investigate EC-SMC interactions and IH development and subjected to arterial flow conditions in a bioreactor. In addition, TEBVs with a confluent endothelial layer underwent POBA-treatment and implantation of bare metal stents (BMS) and drug-eluting stents (DES) to evaluate their suitability as preclinical model for coronary interventions. TEBVs were monitored using optical coherence tomography (OCT).

## Methods

The data that support the findings of this study are available from the corresponding author upon reasonable request. For additional details, please refer to the Major Resources Table (**Table 1**).

**Table 1.**
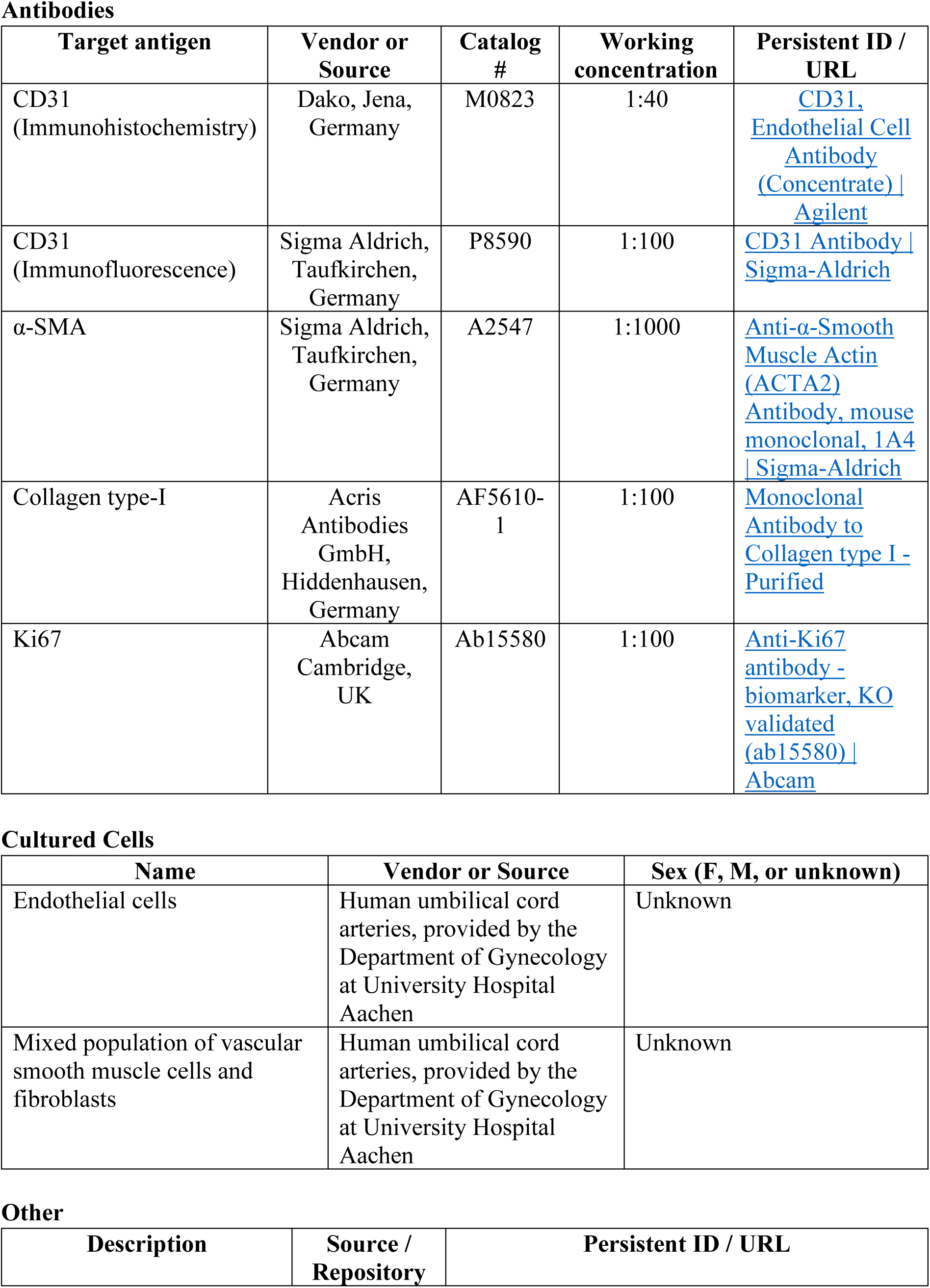
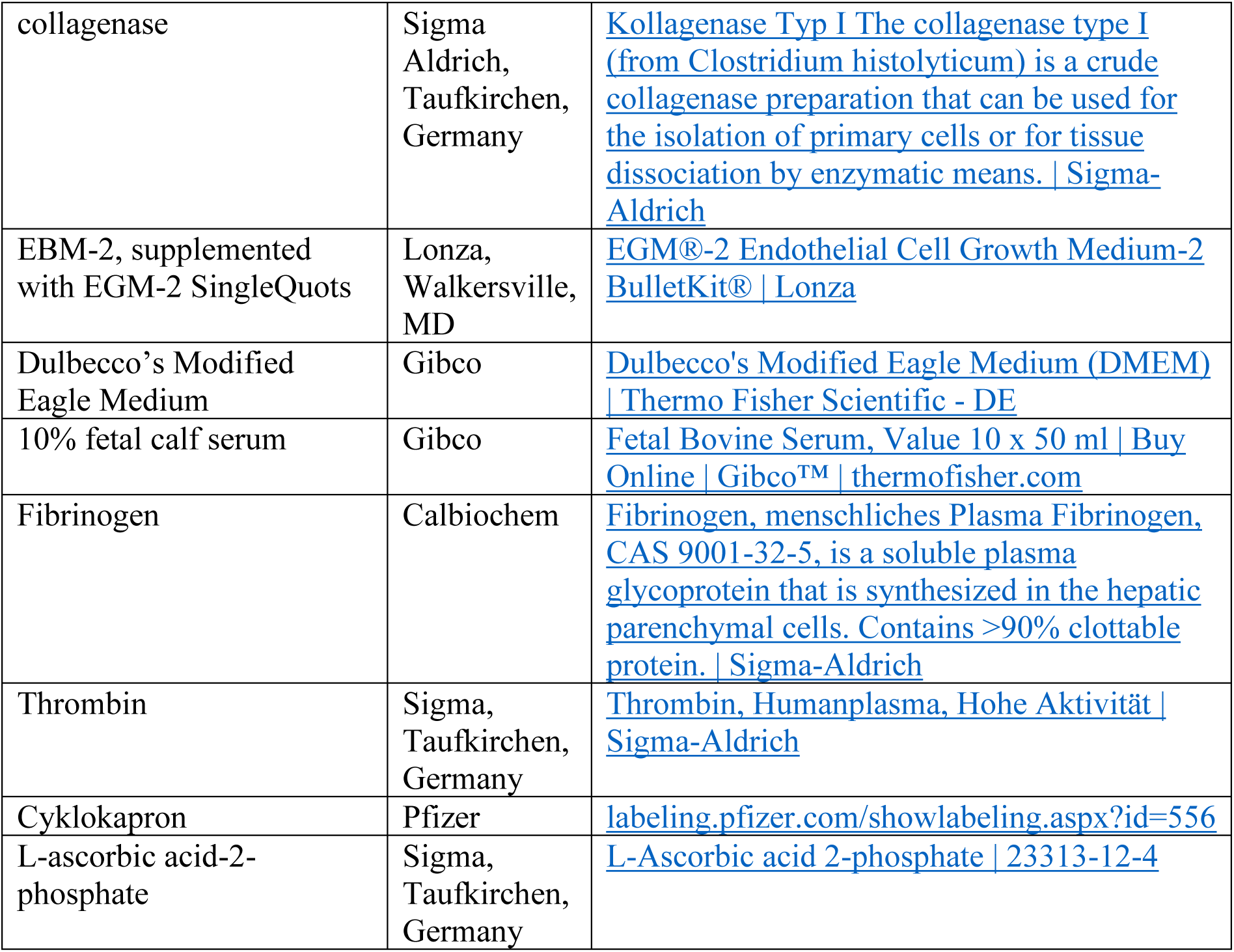
Major Resources Table.

Antibodies
| Target antigen | Vendor or Source | Catalog # | Working concentration | Persistent ID / URL |
| --- | --- | --- | --- | --- |
| CD31<br>(Immunohistochemistry) | Dako, Jena,<br>Germany | M0823 | 1:40 | CD31, Endothelial Cell Antibody (Concentrate) \| Agilent |
| CD31<br>(Immunofluorescence) | Sigma Aldrich,<br>Taufkirchen,<br>Germany | P8590 | 1:100 | CD31 Antibody \| Sigma-Aldrich |
| α-SMA | Sigma Aldrich,<br>Taufkirchen,<br>Germany | A2547 | 1:1000 | Anti-α-Smooth Muscle Actin (ACTA2) Antibody, mouse monoclonal, 1A4 \| Sigma-Aldrich |
| Collagen type-I | Acris<br>Antibodies<br>GmbH,<br>Hiddenhausen,<br>Germany | AF5610-1 | 1:100 | Monoclonal Antibody to Collagen type I - Purified |
| Ki67 | Abcam<br>Cambridge,<br>UK | Ab15580 | 1:100 | Anti-Ki67 antibody - biomarker, KO validated (ab15580) \| Abcam |

Cultured Cells
| Name | Vendor or Source | Sex (F, M, or unknown) |
| --- | --- | --- |
| Endothelial cells | Human umbilical cord arteries, provided by the Department of Gynecology at University Hospital Aachen | Unknown |
| Mixed population of vascular smooth muscle cells and fibroblasts | Human umbilical cord arteries, provided by the Department of Gynecology at University Hospital Aachen | Unknown |

Other
| Description | Source / Repository | Persistent ID / URL |
| --- | --- | --- |
| collagenase | Sigma Aldrich, Taufkirchen, Germany | Kollagenase Typ I The collagenase type I (from Clostridium histolyticum) is a crude collagenase preparation that can be used for the isolation of primary cells or for tissue dissociation by enzymatic means. \| Sigma-Aldrich |
| EBM-2, supplemented with EGM-2 SingleQuots | Lonza, Walkersville, MD | EGM®-2 Endothelial Cell Growth Medium-2 BulletKit® \| Lonza |
| Dulbecco's Modified Eagle Medium | Gibco | Dulbecco's Modified Eagle Medium (DMEM) \| Thermo Fisher Scientific - DE |
| 10% fetal calf serum | Gibco | Fetal Bovine Serum, Value 10 x 50 ml \| Buy Online \| Gibco™ \| thermofisher.com |
| Fibrinogen | Calbiochem | Fibrinogen, menschliches Plasma Fibrinogen, CAS 9001-32-5, is a soluble plasma glycoprotein that is synthesized in the hepatic parenchymal cells. Contains >90% clottable protein. \| Sigma-Aldrich |
| Thrombin | Sigma, Taufkirchen, Germany | Thrombin, Humanplasma, Hohe Aktivität \| Sigma-Aldrich |
| Cyklokapron | Pfizer | labeling.pfizer.com/showlabeling.aspx?id=556 |
| L-ascorbic acid-2-phosphate | Sigma, Taufkirchen, Germany | L-Ascorbic acid 2-phosphate \| 23313-12-4 |

**Table 2.**

| Device | Brand | Company | Diameter<br>(mm) | Strut<br>Thickness<br>(µm) | Nominal<br>Pressure<br>(atm) |
| --- | --- | --- | --- | --- | --- |
| Coronary<br>Dilatation<br>Catheter | TREK™ | Abbott,<br>Chicago, IL | 3.5 | N/A | 8 |
| Bare Metal<br>Stent | Coroflex®<br>Blue Neo | B. Braun | 3.5 | 60 | 10 |
| Drug-Eluting<br>Stent | Promus<br>Element™<br>Plus | Boston<br>Scientific | 3.5 | 81 | 11 |

### Cell isolation and culture

ECs as well as a mixed population of SMCs and FBs were isolated from human umbilical cord arteries provided by the Department of Gynecology at University Hospital Aachen. All human subjects gave their informed consent, and the study protocol was approved by the Independent Ethics Committee at the RWTH Aachen Faculty of Medicine (EK 2067).

After removal of adventitial tissue, arteries were washed with phosphate buffered saline (PBS; Thermo Fisher Scientific), followed by incubation with 1 mg/mL collagenase (Sigma-Aldrich, Taufkirchen, Germany) for enzymatic removal of ECs. Arteries were flushed with EC growth medium (EBM-2, supplemented with EGM-2 SingleQuots, Lonza, Walkersville, MD) to harvest ECs. Isolated ECs were seeded on tissue culture flasks pre-coated with 2% gelatin and incubated with EC growth medium.

In addition, to isolate a mixed population of SMCs and FBs, arteries were cut into tissue rings of 1 mm in length and placed in tissue culture flasks (Corning). Outgrowing cells were observed after 1–2 weeks. SMC/FB mixtures were cultured in Dulbecco’s Modified Eagle Medium (DMEM; Gibco) supplemented with 10% fetal calf serum (FCS; Thermo Fisher Scientific) and subcultured at 80 – 90% of confluence. Cells in passages 2 to 3 for ECs and passages 5 to 7 for SMC/FB mixtures were used to fabricate the TEBVs.

### Fibrin gel

Lyophilized fibrinogen (Calbiochem) was dissolved in Milli-Q purified water (Millipore) and dialyzed overnight against tris-buffered saline (TBS; pH 7.4) using a membrane with a molecular weight cut-off of 6000–8000 (Novodirect). Subsequently, the fibrinogen solution was filter-sterilized, and the concentration was optically determined by measuring the absorbance at 280 nm using an Infinite M200 spectrophotometer (Tecan Group Ltd). Fibrinogen concentration was adjusted to 10 mg/mL by addition of TBS as required.

SMC/FB mixtures were suspended in 1.4 mL TBS at a concentration of 10 × 10^6^ cells /mL. To fabricate one TEBV of 5 cm in length, 4 mL of fibrin gel was prepared from 2 mL fibrinogen solution, 1.4 mL SMC/FB mixtures in TBS, 0.3 mL 50 mM CaCl_2_ (Sigma) in TBS, and 0.3 mL 40 U/mL thrombin (Sigma).

### TEBV fabrication and conditioning

To prepare the TEBV, a molding system was manufactured as described previously (28, 29), consisting of a 6.4 mm inner diameter silicone tube (Carl Roth) and a cylindrical 3 mm inner cylinder fabricated from stainless steel (**Figure 1A, B**). Accordingly, each TEBV had an outer diameter of 6.4 mm and an inner diameter of 3 mm. Both ends of the mold were connected to T-connectors (Fleima Plastic, Wald-Michelbach, Germany), serving as sprue and riser. To provide mechanical support for the fibrin-based TEBV, a PVDF mesh was fixed between the two T-connectors. Fibrin gel components were inserted into the annular space of the mold to initiate the polymerization process. The inner cylinder was removed after polymerization, and the TEBV was maintained in the silicone tube. Subsequently, the lumen was filled with ECs suspended in EC growth medium. TEBVs were constantly rotated around their axis at 1 rpm for 6 h using a modified peristaltic pump (Ismatec, Wertheim, Germany) to uniformly distribute ECs on the inner lumen surface. The mold containing the TEBV was then connected to the VascuTrainer bioreactor system for conditioning, as described previously (28) (**Figure 1C**). Briefly, the VascuTrainer consisted of a plastic medium reservoir (Thermo Scientific) and silicone tubing (Ismatec), and connected to a small centrifugal pump (702-6882, RS components), which was controlled by a custom-developed control-unit (**Figure 1D**). All TEBVs were maintained at 37 °C and 5% CO_2_ during cultivation.

**Figure 1.**
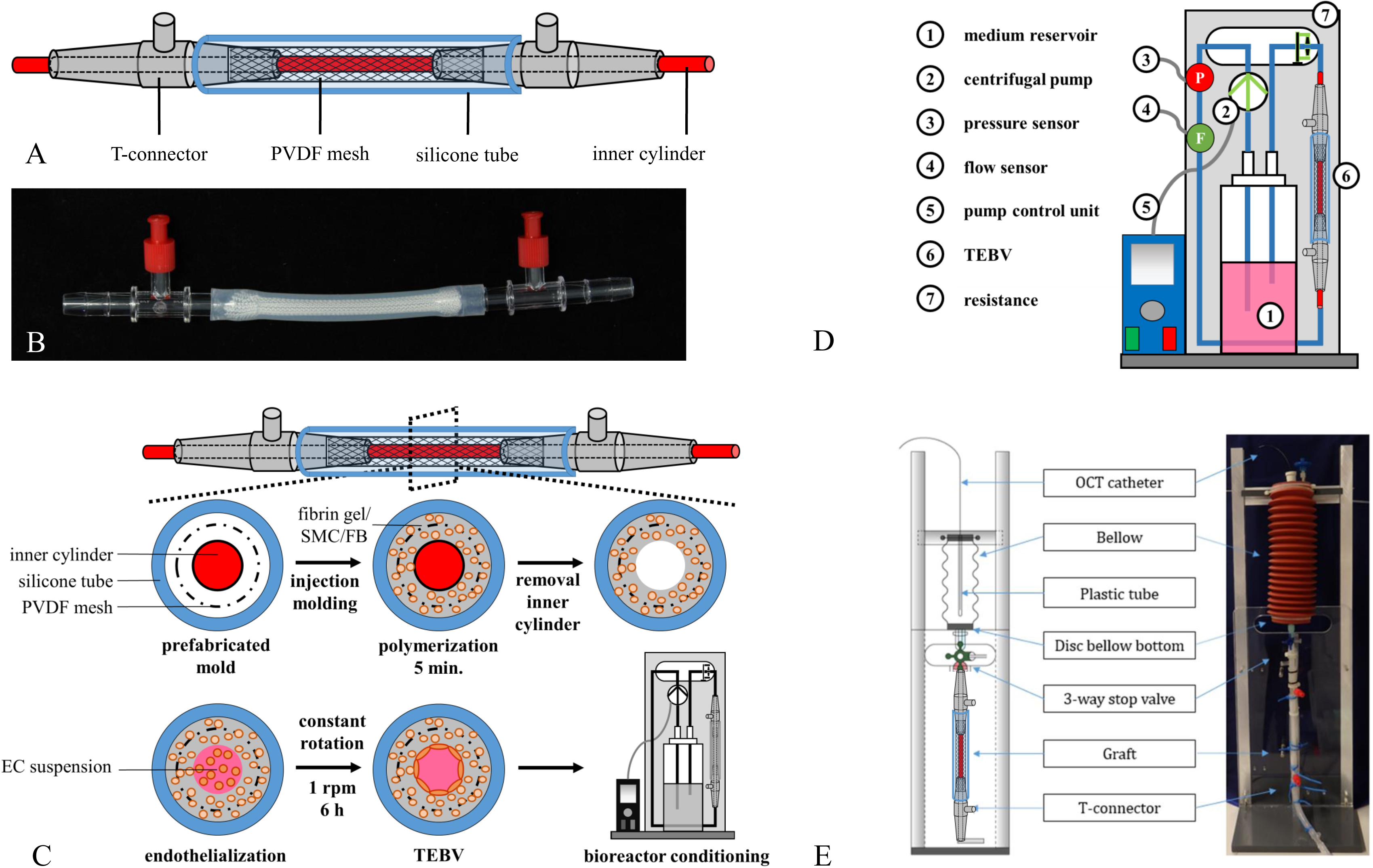
Experimental Setup. **A, B)** Graphic illustrations and photographs of the molding system. **C)** TEBV fabrication. SMCs/FBs in a fibrinogen-CaCl_2_-thrombin mixture were cast into the mold for gel polymerization. After removal of the inner cylinder the lumen was filled with ECs suspended in culture medium, and TEBVs were constantly rotated at 1 rpm for 6 h to uniformly distribute ECs on the inner lumen surface. TEBVs were transferred into the VascuTrainer bioreactor system for conditioning. **D)** Graphic illustration of the VascuTrainer bioreactor system which was assembled from a medium reservoir and silicone tubing and connected to a small centrifugal pump. **E)** A sterile module was implemented to allow for sterile introduction of balloon catheters, stents and catheters for OCT.

In addition, to allow for sterile introduction of balloon catheters, stents, and catheters for OCT, a sterile module was implemented, consisting of a sterile plastic cavity that was accessed through a disinfectable three-way valve (**Figure 1E**). The sterile module was connected to the TEBV-containing mold via a three-way stopcock. Two metal discs were used to seal the cavity at both ends and to maintain sterility. A central hole with a diameter of 2 mm within these discs enabled the introduction and centering of various catheters.

Culture medium in the bioreactor system consisted of a 50:50 (v/v) mixture of DMEM and EGM-2 supplemented with 10% FCS, 1% antibiotic-antimycotic solution (ABM, Gibco), 1.6 µl/mL tranexamic acid (TA) (Cyklokapron® injection solution 1000 mg/mL; Pfizer Pharma GmbH), and L-ascorbic acid-2-phosphate (1.0 mM; Sigma). TEBVs were conditioned for 14 days under pulsatile flow (70 bpm) at arterial levels of shear stress (10 dynes/cm²) and pressure (80–120 mmHg).

### Endothelialization conditions

In order to test the effect of incomplete endothelialization on IH, three different scenarios were created and evaluated (**Figure 2**): (A) TEBV with a confluent EC luminal layer where ECs were seeded at a concentration of 3 × 10^6^ cells/mL, (B) TEBV with a semi-confluent endothelial layer with ECs seeded at a concentration of 0.5 × 10^6^ cells/mL, and (C) TEBV without an EC layer. All experiments were performed in triplicates. EC confluence / semi-confluence was confirmed by immunohistochemical staining against CD31.

**Figure 2.**
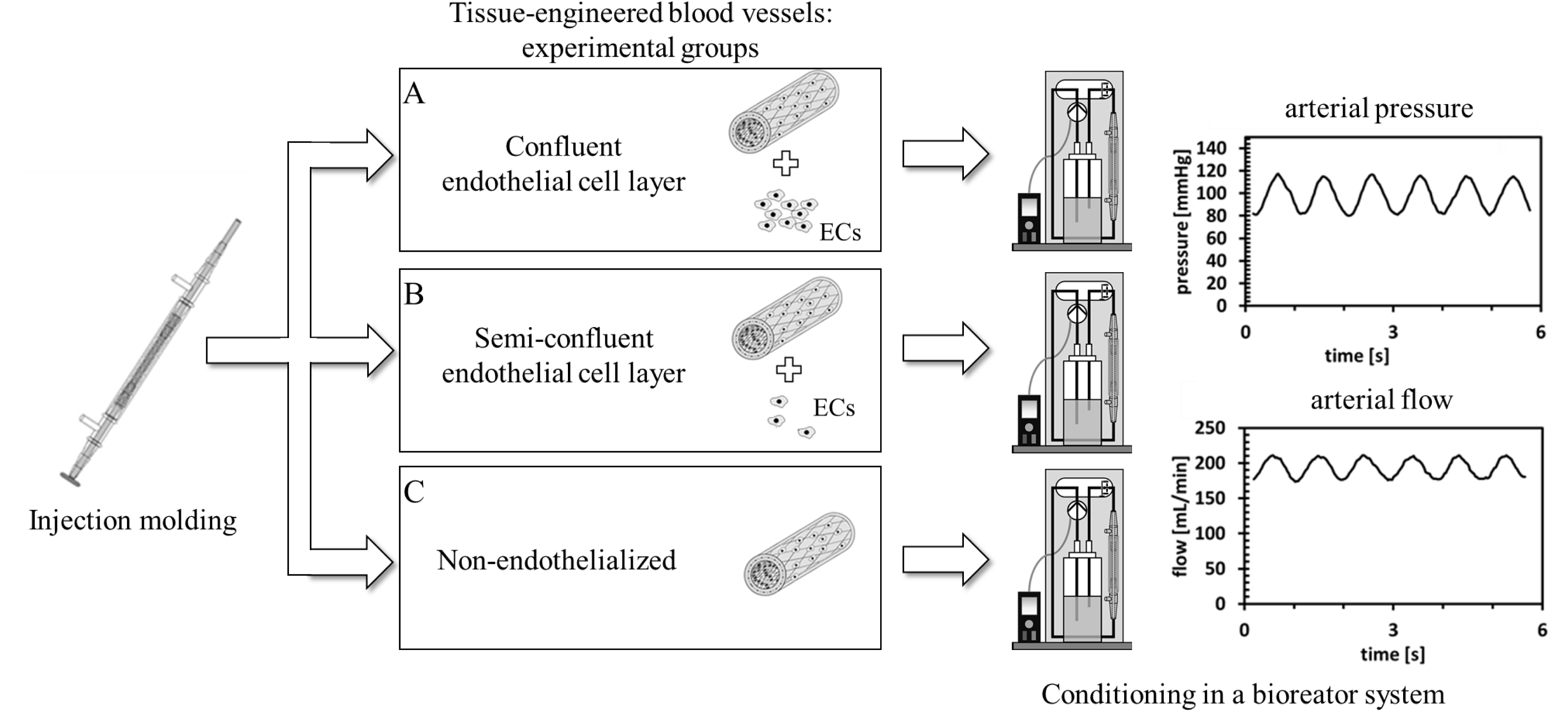
Experimental Conditions. To explore the impact of the endothelial layer on intimal hyperplasia, we compared three different endothelialization conditions of TEBVs. **A)** ECs were seeded at a concentration of 3× 10^6^ cells/mL to generate a confluent EC luminal layer. **B)** To simulate a semi-confluent endothelium, TEBV were fabricated with ECs seeded at a concentration of 0.5 × 10^6^ cells/mL. **C)** TEBV were conditioned without an EC layer. All vessels were subjected to the same arterial pressure and flow conditions.

### Balloon angioplasty and stent implantation

Plain old balloon angioplasty (POBA) and stent implantation were performed in TEBVs with confluent EC layers 7 days after fabrication. For POBA, 3.5 mm TREK balloons (Abbott, Chicago, IL) were used. Stent implantation was performed using balloon-mounted bare metal stents (BMS; Coroflex Blue Neo, B. Braun, Melsungen, Germany) and drug-eluting stents (DES; Promus Element Plus, Boston Scientific). All experiments were performed in triplicates.

Using a guidewire (HI-TORQUE Balance Middle Weigth Universal Guide Wire, Abbott), a plain angioplasty balloon or a balloon-mounted human coronary stent was introduced into the lumen of the TEBV and placed at 1 cm below the connector end used as point of access. Balloon inflation and stent deployment were performed at defined nominal pressures for each device (**Table 1**), using an insufflator (Encore™, Boston Scientific). Untreated TEBVs with a confluent EC layer served as control. Following POBA or stent implantation, TEBVs were conditioned for additional 14 days (*i.e.*, 21 days in total).

### Imaging

Non-invasive external ultrasound monitoring of TEBV integrity and visualization of stent position over the course of the conditioning period was performed using a portable ultrasound system, Vivid-I (S/N 3642VI, General Electric), equipped with a linear probe (8L-RS, General Electric).

Intravascular imaging with OCT to provide cross-sectional and three-dimensional images of the TEBVs was performed after 14 days of conditioning using the Dragonfly OpStar™ Imaging Catheter with the OPTIS™ Imaging System (both Abbott).

Macroscopic images of the cross-section of the TEBVs with different degrees of endothelialization, were captured using a digital camera (Nikon Corporation, Japan). Each image was analyzed using Fiji (ImageJ) (30), and wall thickness was calculated as the mean of 12 measurements taken at different positions around the graft circumference for each sample.

### Histological and immunohistochemical analysis

TEBVs cultured under different endothelialization conditions were immersion fixed in Carnoy’s solution, dehydrated in increasing series of ethanol and embedded in paraffin for sectioning. For immunohistochemistry, paraffin sections were deparaffinized before antigen retrieval by steam heat in citrate buffer (pH 6.0). The sections were treated with 3% H_2_O_2_. Non-specific antibody binding sites were blocked using Dako protein block (Dako, Jena, Germany) for 10 min. before incubation with a primary antibody against CD31 (mouse monoclonal anti-CD31 (1:40; Dako)). Antibody binding was detected using a streptavidin-HRP solution (Dako).

Stented and POBA-treated TEBVs, as well as their untreated control TEBVs, were fixed with 4% formalin (Carl Roth, Karlsruhe, Germany) and embedded in polymethyl-methacrylate (PMMA; Technovit 9100, Kulzer, Wertheim, Germany). Sequential sections from the proximal, mid, and distal parts of the stented TEBVs were sawed and grinded to sections of 40 – 50 µm thickness using EXACT thin section cutting system (EXACT, Norderstedt, Germany). Sections of all TEBVs were stained with toluidine blue/ basic fuchsine solution, staining nuclei dark blue and matrix proteins red, according to the manufacturer’s instructions.

Images were obtained from cross-sections using a DMI-3000 microscope (Leica, Wetzlar, Germany) and DISKUS image analysis software (version 4.80, C. Hilgers Technical Bureau, Königswinter, Germany). Measurement of lumen and wall thickness was performed by tracing the contours of the outer TEBV wall and of the lumen area using a Tavla pen tablet (Braun Photo Technik, Nuremberg, Germany). In stented TEBVs, neointimal growth was measured above the stent struts.

### Immunofluorescence staining

Paraffin-embedded TEBVs were deparaffinized, non-specific binding sites were blocked and the cells were permeabilized by incubation in 5% normal goat serum (NGS, Dako) in 0.1% Triton-X-100 (Sigma) in PBS. Sections were incubated for 1h at 37°C with the following primary antibodies: mouse monoclonal anti-α-SMA (1:1000; Sigma), rabbit anti-type-I collagen (1:100; Acris Antibodies GmbH, Hiddenhausen, Germany), mouse monoclonal anti-CD31 (1:100; Sigma), and rabbit monoclonal anti-Ki67 (1:100; Abcam Cambridge, UK). The sections were then incubated for 1 h at room temperature with either rhodamine-or fluorescein-conjugated secondary antibodies (1:400; Molecular Probes, Leiden, Netherlands). Cell nuclei were counterstained using DAPI nucleic acid stain (Molecular Probes). As negative controls, samples were incubated in diluent and the secondary antibody only, and samples from umbilical cord artery served as positive controls. Images were acquired with a fluorescence microscope (AxioObserver Z1; Carl Zeiss GmbH, Oberkochen, Germany), equipped with a digital color camera (AxioCam MRm; Carl Zeiss GmbH,).

### Collagen content quantification

Collagen levels were determined as described before (31) by measuring the hydroxyproline content of overnight vacuum-dried samples from each TEBV. Briefly, samples were hydrolyzed in 500 µL of 6M hydrochloric acid (HCL) at 110°C for 18 h. Buffered chloramine-T reagent (60 mM, 20% n-propanol and 80% acetate citrate buffer) was added to oxidize the samples for 45 min. Subsequently, samples were incubated with Ehrlich’s reagent (4-dimethylamino.benzaldehyde, Sigma) at 65°C for 45 min. Absorbance was measured at 550 nm using an Infinite M200 spectrophotometer (Tecan, Männedorf, Switzerland). A standard curve was generated using known amounts of trans-4-hydroxy-L-proline (Sigma), and hydroxyproline concentration was calculated for each sample.

### Cytokine measurements

Medium samples for cytokine analysis were collected from the bioreactor at 0.5, 1, 2, and 24 hours, and at 14 days (end of conditioning) after POBA or stent implantation. Medium samples from untreated fully endothelialized TEBVs served as controls and were obtained simultaneously. We used DuoSet ELISA kits (R&D Systems, Minneapolis, USA) to assess levels of interleukin 6 (IL-6), interleukin 8 (IL-8), and monocyte chemoattractant factor 1 (MCP-1).

### Statistical analysis

Statistical analyses and creation of graphs were performed with GraphPad Prism (GraphPad Software, Version 10.0.2.232). Continuous data are presented as mean ± SD. We used Shapiro-Wilk test to assess normality. Statistical differences were determined with Student’s t test and one-way ANOVA, followed by Tukey multiple comparisons testing for normally distributed variables, and with Mann-Whitney U test and Kruskal-Wallis test followed by Dunn’s multiple comparisons testing for non-normally distributed variables.

## Results

### Impact of endothelial integrity on wall thickening

To test the effect of an incomplete endothelial layer on vascular wall thickening and IH, TEBVs were generated with a confluent EC luminal layer, with a semi-confluent endothelium, and without an EC layer. After 14 days of dynamic conditioning under arterial flow and pressure conditions, TEBVs underwent OCT-imaging. Introduction of the OCT-catheter and imaging was feasible in all TEBVs. TEBVs with absent or incomplete endothelialization exhibited a reduced lumen diameter in OCT compared with TEBVs generated with a confluent EC layer (**Figure 3A**).

**Figure 3.**
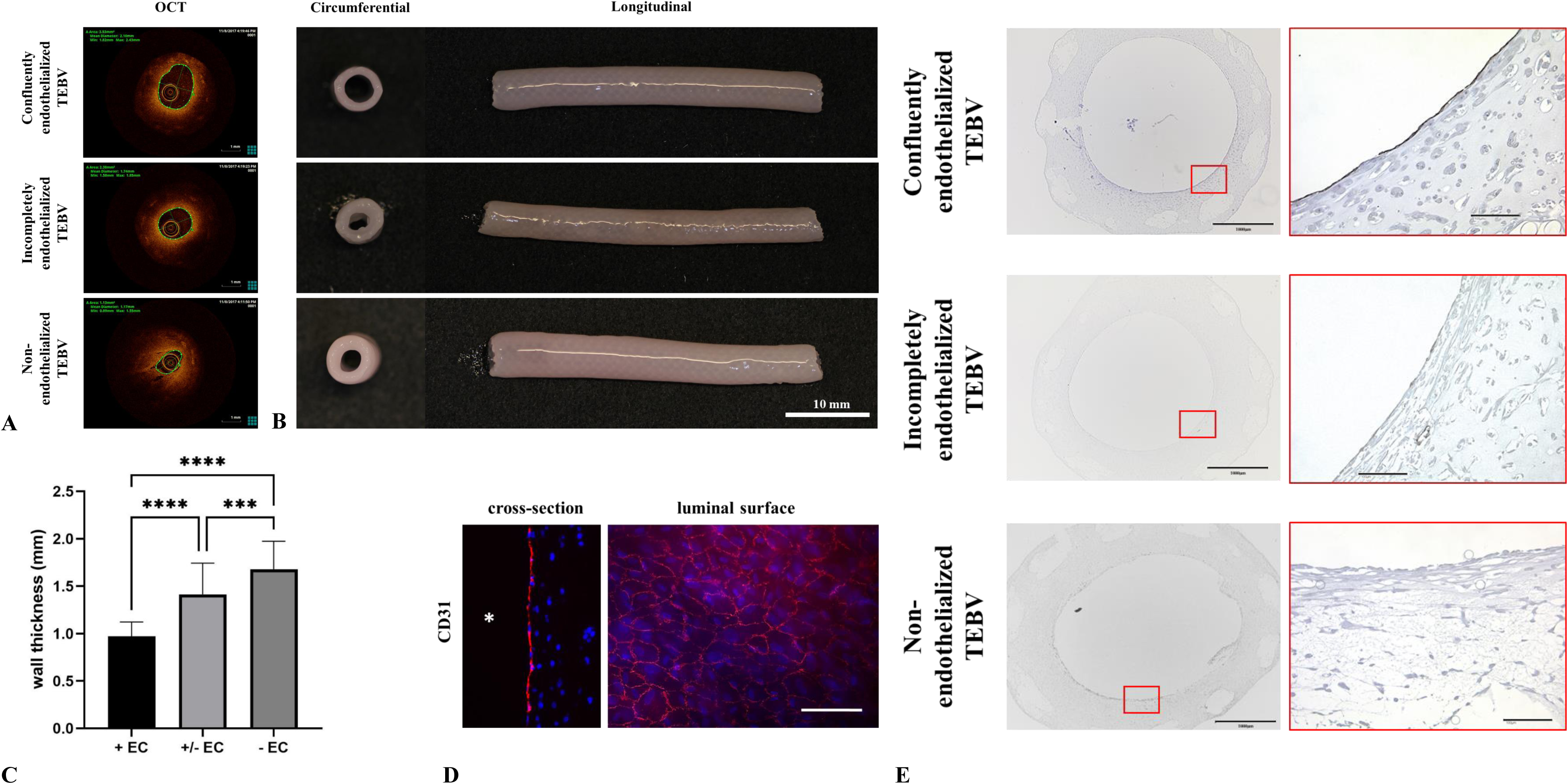
Impact of endothelial integrity on wall thickening. **A, B)** TEBVs with absent or incomplete endothelialization exhibited a reduced lumen diameter compared with TEBVs generated with a confluent EC layer, as verified by OCT **(A)** and macroscopically **(B)**. **C)** Compared with TEBVs that were confluently seeded with ECs, TEBVs with absent or incomplete endothelial layer exhibited significantly thicker vessel walls (****P<0.0001; ***P<0.005). **D, E)** Immunofluorescence **(D)** and immunohistochemical **(E**, upper panel) staining against CD31 confirmed a confluent EC layer on the inner surface of TEBVs seeded with ECs at a concentration of 3 × 10^6^ cells/mL. Endothelialization was incomplete in TEBVs that were seeded with ECs at lower cell concentration **(E**, middle panel). No CD31-positive cells were observed in TEBVs without ECs (**E**, lower panel). Scale bars: 1000µm (overview, left), 100µm (cut-out, right).

In line with OCT-imaging results, macroscopic analysis of TEBVs without ECs showed significantly thicker vessel walls (1.68 ± 0.30mm) compared with TEBVs with an incomplete (1.41 ± 0.33mm) and with a confluent EC luminal layer (0.97 ± 0.15mm) (**Figure 3B, C**). While immunofluorescence and immunohistochemical staining against CD31 confirmed evenly distributed ECs along the inner surface of the TEBV wall (**Figure 3D, E**), endothelialization was incomplete in TEBVs that were seeded with ECs at lower cell concentration (**Figure 3E**).

### Cellular composition and proliferation in TEBVs with intact, semi-confluent, and absent endothelium

Cellular proliferation and composition of TEBV vessel walls were assessed by immunofluorescence microscopy (**Figures 4 and 5**). We performed immunofluorescence staining against αSMA, a cytoskeleton protein mostly synthetized by vascular SMCs and SMC-like cells such as myofibroblasts which is responsible for SMC contractile properties (32, 33). While a substantial number of cells stained positive for αSMA within TEBV vessel walls with confluent or semi-confluent endothelial lining, αSMA-positive cells were sparse in TEBVs without ECs (**Figure 4**).

**Figure 4.**
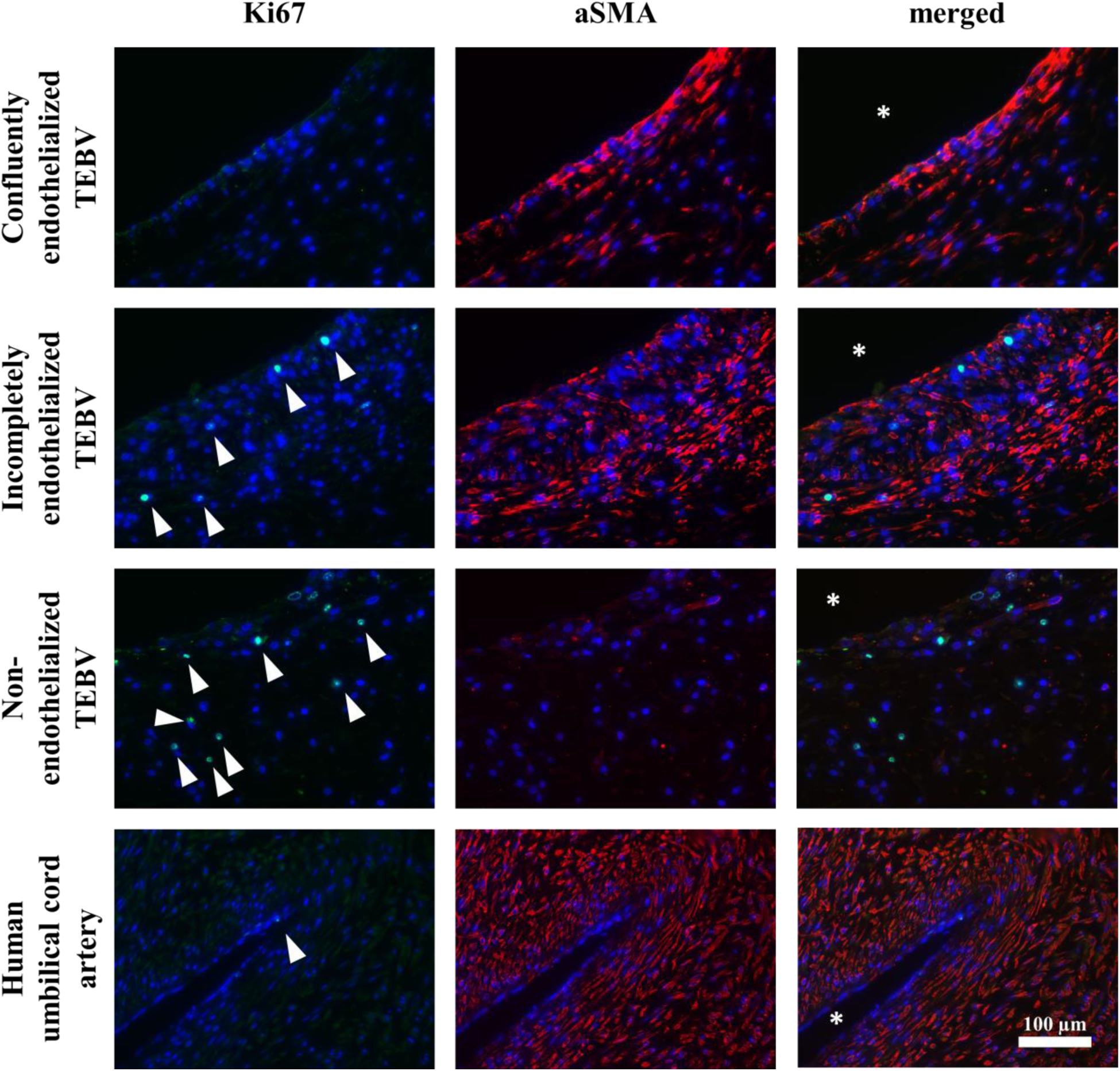
Cellular proliferation and composition of TEBV vessel walls. In confluently or partially endothelialized TEBVs, a substantial number of cells stained positive for αSMA, indicating contractile properties of SMCs. However, αSMA-positive cells were sparse in TEBVs without ECs. Staining against Ki67 revealed more proliferating cells (white arrowheads) in TEBVs without ECs than in those with a semi-confluent or intact endothelial layer. Human umbilical cord served as control * indicates lumen area.

**Figure 5.**
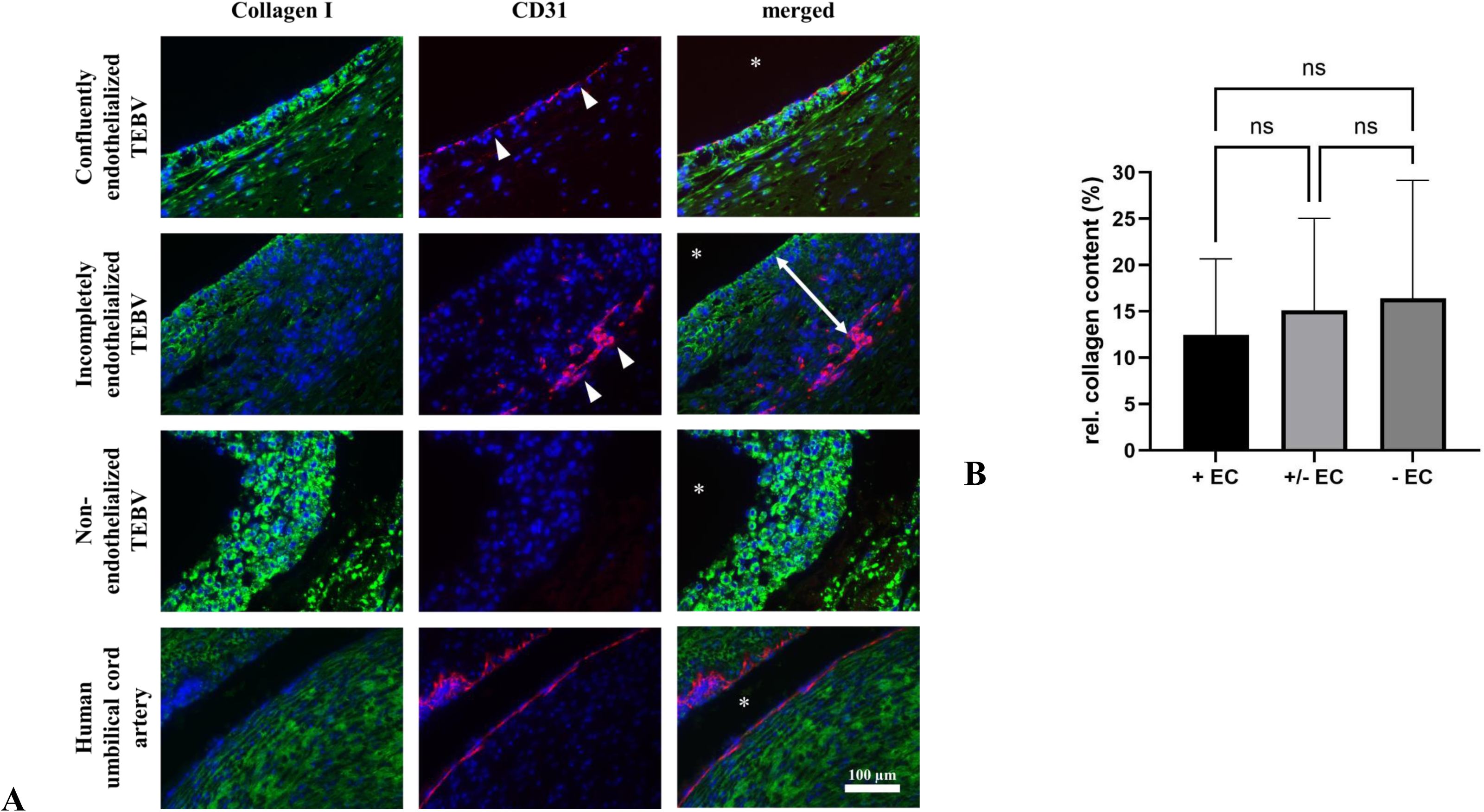
Endothelialization and collagen content in TEBVs. **A)** Immunofluorescence double staining against collagen I and CD31. In TEBVs with a confluent endothelial layer, collagen appeared organized in elongated fibers, and CD31-positive ECs (white arrowheads) formed a continuous, uniform layer on the luminal surface. In TEBVs seeded with ECs at lower cell concentration, CD31-positive cells were found within the TEBV wall, covered by a layer of CD31-negative cells, and collagen was detectable primarily at the luminal side of the vessel wall. Collagen-I staining was most pronounced in non-endothelialized TEBVs, although a clear fibrillar structure was not observed. **B)** Relative to native arteries (50µg/mg), collagen content, assessed by hydroxyproline assay, was highest in TEBVs without ECs and lowest in confluently endothelialized TEBVs, with incompletely endothelialized TEBVs in between, however, these differences were not statistically significant (ns = not significant).

To assess cellular proliferation in TEBVs, we performed immunofluorescence staining against Ki67. We observed a greater presence of cells in a proliferative state in TEBVs without ECs compared with TEBVs with semi-confluent and intact endothelium (**Figure 4**), suggesting enhanced proliferation in the absence of EC-mediated regulation.

Taken together, double staining against αSMA and Ki67 suggested an inverse relationship between the synthesis of αSMA and cell proliferation.

### Collagen content in TEBVs with intact, semi-confluent, and absent endothelium

Next, to localize and quantify the extent of matrix protein deposition, we performed immunofluorescence staining against collagen I and CD31 (**Figure 5A**). Interestingly, we found CD31-positive cells within the TEBV wall, covered by a layer of CD31-negative cells in TEBVs seeded with ECs at lower cell concentration. Collagen-I staining appeared more extensive in non-endothelialized TEBVs compared with partially and completely endothelialized samples. In TEBVs with a confluent endothelial layer, collagen appeared organized in elongated fibers. In partially endothelialized TEBVs, such organization was detectable primarily at the luminal side of the vessel wall, whereas non-endothelialized TEBVs exhibited no clear fiber structure.

For quantitative assessment of collagen content in TEBVs, hydroxyproline concentration, a major amino acid constituent of collagen, was measured in samples of TEBVs. In line with our findings from immunofluorescence staining, mean hydroxyproline content was highest in TEBVs without ECs while it was lowest in TEBVs with an intact endothelial layer, albeit without statistical significance (**Figure 5B)**. Relative to the average amount of hydroxyproline found in native arteries (50µg/mg) (34), hydroxyproline content was 12.49 ± 8.20% in TEBVs with an intact endothelial layer, 15.12 ± 9.95% in TEBVs with incomplete endothelialization, and 16.41 ± 12.75% in TEBVs without ECs.

### Plain-old balloon dilatation and stent implantation of TEBVs

To assess whether TEBVs could serve as *in vitro* models for the evaluation of coronary devices, we performed POBA and stent implantation (**Figure 6A**) in TEBVs at 7 days of conditioning. Non-treated TEBVs seeded with a confluent layer of ECs served as controls (all n=3). All procedures were carried out successfully, with adequate trackability of balloons and balloon-mounted coronary stents over a guidewire. No TEBVs were obviously damaged during the procedure. Over the course of the conditioning period of 14 days after the procedures (*i.e*., 21 days in total), no infection occurred, and all bioreactors remained sterile. Ultrasound monitoring was carried out in all TEBVs and proved useful for non-invasive monitoring as boundaries of both the stents and of TEBV walls were clearly definable (**Figure 6B-E**). TEBVs remained patent throughout the observation period, and stents maintained their original position. In addition, at 14 days of conditioning after POBA and stent implantation, all TEBVs underwent OCT-imaging (**Figure 6F, G**). Incipient cellular growth was visible above the stent struts as small reflection points. Following OCT-imaging, TEBVs were removed from the bioreactor for macroscopic inspection (**Figure 6H-K**).

**Figure 6.**
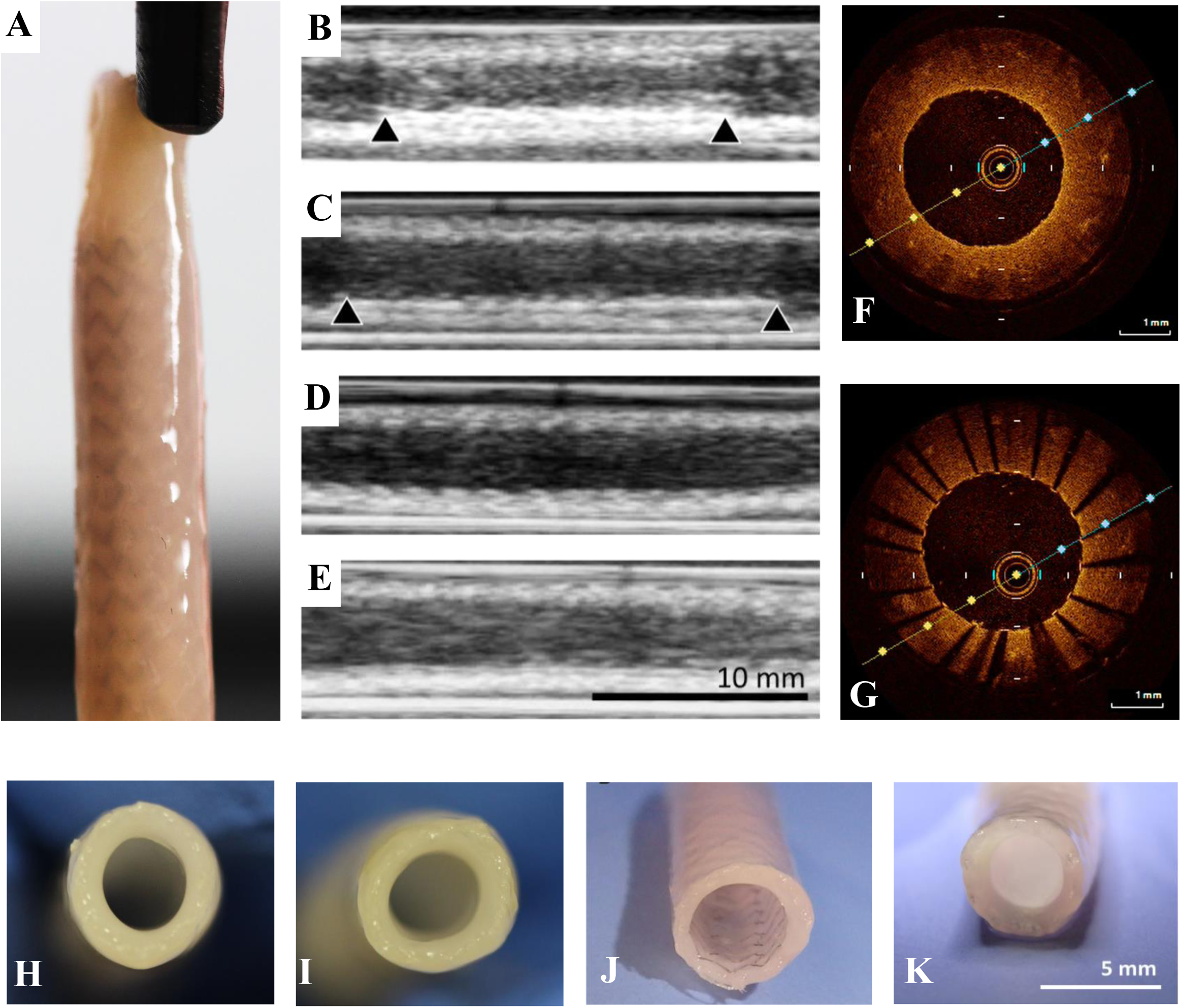
POBA and stent implantation in TEBVs. **A)** TEBV with a commercially available human-sized coronary stent. **B-E)** Ultrasound monitoring proved useful for non-invasive monitoring after implantation of BMS **(B)** and DES **(C)**, after POBA **(D)**, and in untreated control TEBVs **(E)**. Boundaries of the stents (black arrowheads) as well as the TEBV vessel walls were clearly visible. **F, G)** OCT-imaging was performed 14 days after POBA and stent implantation. **H-K)** Macroscopic views of control **(H)**, POBA-treated **(I)**, BMS-treated **(J)**, and DES-treated **(K)** TEBVs.

### Histology of TEBVs after POBA and stent implantation

TEBVs were embedded in PMMA and processed for histology (**Figure 7A-D)**. Vessel wall thickness was highest in POBA-treated TEBVs (0.8mm ± 0.1mm) compared with untreated TEBV controls (0.64mm ± 0.11mm) and compared with BMS-(0.59mm ± 0.09mm) and DES-(0.63mm ± 0.13mm) treated TEBVs (p<0.001, **Figure 7E**). There were no significant differences in vessel wall thickness among other comparison groups. In particular, wall thickness was comparable between BMS- and DES-treated TEBVs.

**Figure 7.**
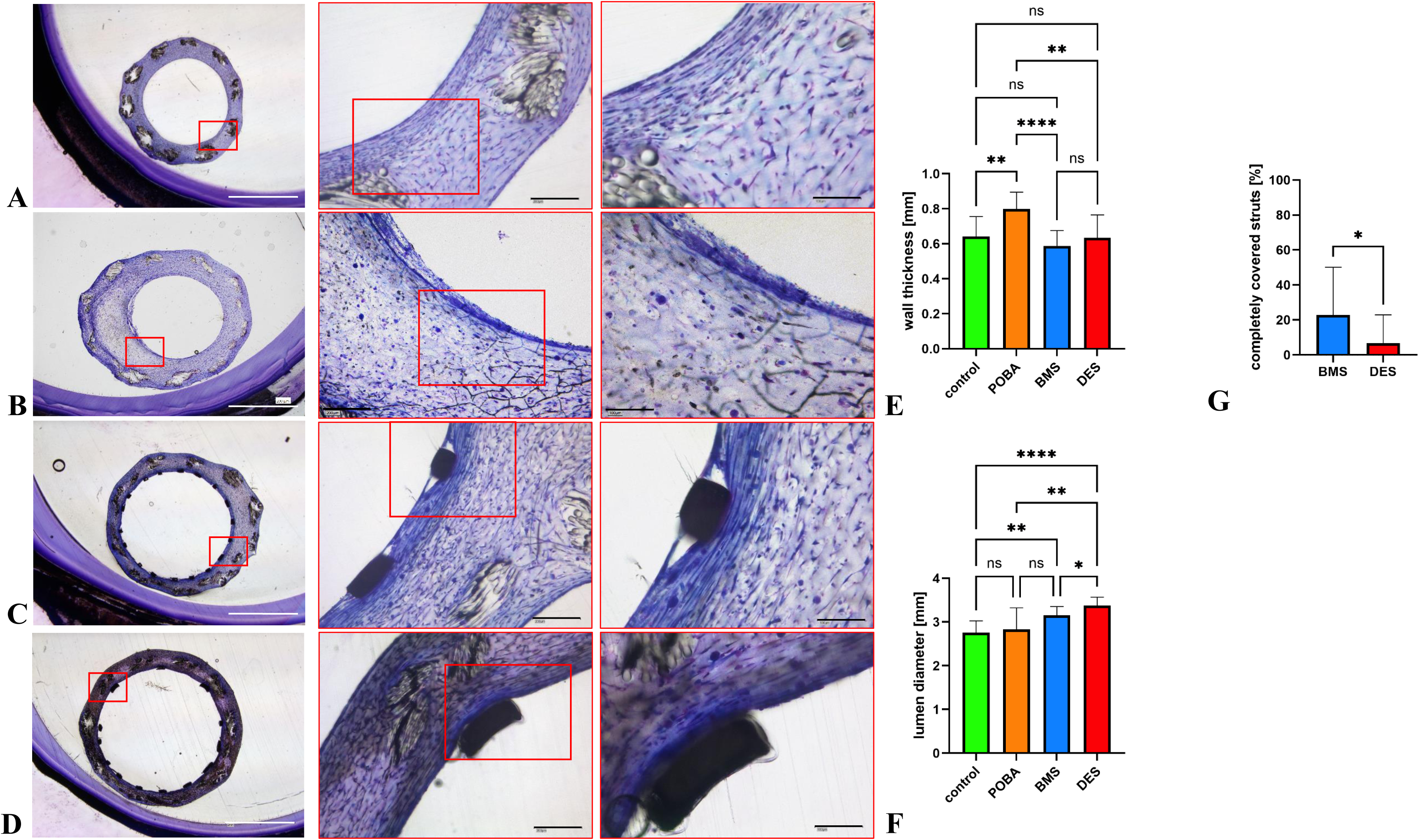
Histology of stented and POBA-treated TEBVs compared with untreated control TEBVs. **A-D)** Untreated control **(A),** POBA- **(B)**, BMS- **(C)**, and DES- **(D)** treated TEBVs were embedded in plastic and processed for histology (toluidine blue/ fuchsine staining). **E)** Vessel wall thickness was higher in POBA-treated TEBVs compared with untreated TEBV controls and stented TEBVs. **F)** Lumen diameter was greater in stented compared with POBA-treated and untreated control TEBVs. DES-treated TEBVs exhibited the largest lumen diameter. No significant difference in lumen diameter was observed between BMS- and POBA-treated TEBVs, or between POBA-treated TEBVs and untreated controls. **G)** 14 days after stent implantation, 22.9% of struts were completely covered with neointima in BMS-treated TEBVs, while it were only 6.8% of struts in DES-treated TEBVs. ****P<0.0001; ***P<0.005, **P<0.01, *P<0.05.

Lumen diameter was greater in stented TEBVs compared with POBA-treated (2.83mm ± 0.49mm) and untreated control TEBVs (2.75mm ± 0.27mm), with DES-treated TEBVs exhibiting the largest lumen (3.38mm ± 0.19mm), which was also significantly larger compared with BMS-treated TEBVs (3.15mm ± 0.21mm, **Figure 7F)**. No significant difference was observed between BMS- and POBA-treated TEBVs, or between POBA-treated and untreated controls.

We found incipient neointima formation around the stent struts within TEBVs treated with BMS (**Figure 7C**). In contrast, there was only marginal neointimal growth within TEBVs treated with DES (**Figure 7D**). In cross-sections of BMS-treated TEBVs, 22.9% of struts were completely covered with neointima, while it was only 6.8% of struts in DES-treated TEBVs (**Figure 7G**).

### Inflammatory cytokine levels following POBA and stent implantation

Over the course of the first 24h of the conditioning period after stent implantation, we observed a steady increase of levels of IL-6, IL-8, and MCP-1 in culture medium samples from stented and untreated control TEBVs (**Figure 8**). Cytokine levels at 14 days were significantly higher compared to baseline levels in all samples. The highest levels of IL-6, IL-8, and MCP-1 in culture medium were obtained from bioreactors with BMS-treated TEBVs. There were no significant differences in IL-6-, IL-8-, and MCP-1-levels between DES-treated TEBVs and untreated controls at 14 days.

**Figure 8.**
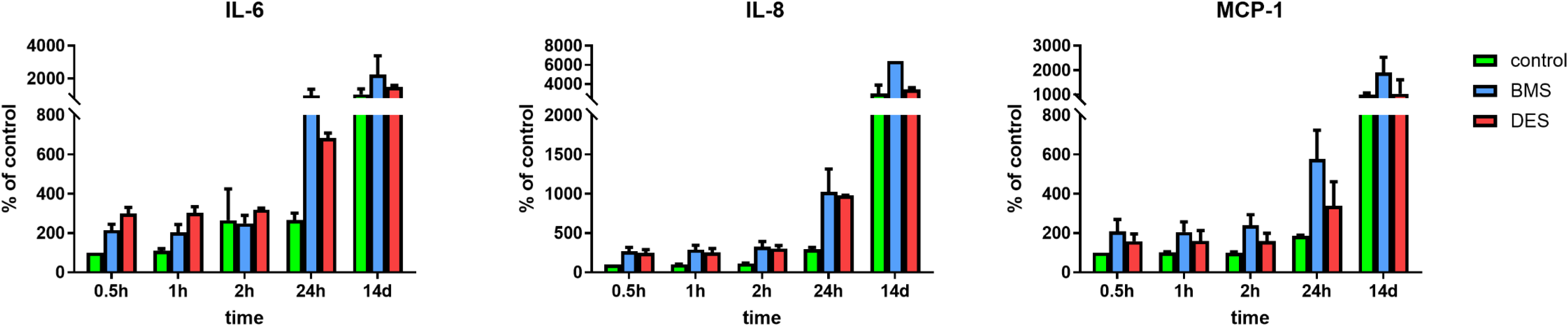
Cytokine levels in culture medium samples from stented vs. untreated control TEBVs. IL-6, IL-8, and MCP-1 levels increased over time in culture medium from stented and control TEBVs, with highest levels in BMS-treated TEBVs.

## Discussion

Here, we present TEBVs as an *in vitro* model for IH and restenosis following POBA and stent implantation. Owing to its tubular shape and its dimensions similar to a human coronary artery, our model not only enables the investigation of EC-SMC crosstalk under shear stress and under different culture conditions, but also allows for POBA and for the implantation of human-sized coronary stents.

EC-SMC crosstalk is crucial to maintain vascular homeostasis. An intact EC layer is essential for the integrity of the vessel wall and contributes to the quiescent state of the underlying SMCs (4, 8–10). ECs also inhibit SMC type I collagen synthesis when co-cultured as a bilayer (9). *Vice versa*, perturbations in the EC-SMC crosstalk can elicit endothelial dysfunction or phenotypic switching of SMCs from a contractile and quiescent towards a synthetically active, migratory and proliferative phenotype (4).

Three-dimensional co-culture models have the potential to replicate not only the anatomical macrostructure but also the microstructure at tissue-and at cell-level, including ECM deposition and arrangement. Therefore, three-dimensional models such as EC-SMC spheroids with direct cultures of ECs and SMCs (35, 36), vascular organoids created from induced pluripotent stems cells (37, 38), and cultures of ECs on ECM-like gels containing SMCs (39) have increased physiological relevance and may bridge the gap between traditional 2D cell culture and *in vivo* animal models (25). However, these models are not able to recapitulate the hemodynamic environment of blood vessels. Under static conditions, ECs induce SMC migration in a co-culture system by increased endothelial expression of platelet-derived growth factor (40–42). Shear stress, however, induces the release of nitric oxide in ECs, which inhibits SMC migration (43–45). Human blood vessels are continuously exposed to mechanical stimuli such as blood pressure and shear stress, and sustained exposure of the human vascular wall to non-physiological flow and pressure conditions may elicit changes in the proliferative and migratory status of cells within the vessel wall, eventually leading to vascular remodeling (6, 46, 47). Therefore, we generated TEBVs as dynamic 3D models which can be exposed to mechanical stimuli. TEBVs in our model were conditioned in a bioreactor ensuring a continuous exposure to physiological flow and pressure conditions.

### TEBVs as *in vitro* models for the research on atherosclerosis

The introduction of triple-layered TEBVs in 1998 (48) laid the foundation for tremendous research activities to further refine TEBV development with both autologous (49) and allogenic cells (50, 51). Originally intended for the use as vascular grafts for coronary artery bypass surgery, for the treatment of peripheral artery disease, or for hemodialysis access (51, 52), TEBVs have also been evaluated as models for *in vitro* atherosclerosis research (53), and as testing platforms for vascular interventions such as POBA and stent implantation (54–56).

More than 20 years ago, TEBVs with a contractile vascular media were developed from sheets of human SMCs wrapped around a styrene tubular support and displayed contraction and relaxation in response to vasoconstrictor and vasodilator agents (57). Later on, Schmidt et al. presented TEBVs generated from myofibroblasts seeded on biodegradable polyglycolic-acid scaffolds and lined with ECs that were exposed to flow conditions (34). A decade ago, the group suggested their constructs as a model to study essential pathogenic phenomena of atherosclerosis *in vitro* (53). They were able to demonstrate enhanced adhesion and transmigration of monocytes injected into the medium upon endothelial stimulation with either tumor necrosis factor α or low-density lipoprotein cholesterol (LDL-C). This model represented a significant step towards a relevant *in vitro* platform for the systematic assessment of atherosclerotic pathways. However, with hydroxyproline content at only 1% of that in native arteries (34), the model exhibits lower collagen levels and only partially reflects the composition of human coronary arteries. Furthermore, the model was equipped with a tubular backbone scaffold of PGA for mechanical support, which introduces mechanical stiffness of the vessel wall and reduces vascular compliance.

We therefore opted for a fibrin gel matrix, supported by a macroporous PVDF mesh. Fibrin gel not only enables a uniform cell distribution but has also been shown to promote the synthesis of matrix proteins by seeded cells, including elastin and collagen (58–60). As we have demonstrated before, the PVDF mesh in our TEBVs provided adequate mechanical support without limiting the resilience and compliance of the vessel wall (61, 62).

We were able to recapitulate fundamental principles of vascular biology. TEBVs with an absent or incomplete endothelial layer had thicker vascular walls and exhibited a reduced lumen diameter. Moreover, we observed a higher number of proliferating cells as well as an increase in collagen content in TEBVs without ECs and in TEBVs with incomplete endothelial layer. In TEBVs with incomplete endothelialization, CD31-positive ECs did not mark the inner lining but were found within the vascular wall, covered by newly-formed tissue consisting of collagen fibers and of cells that stained negative for αSMA, a marker of a contractile SMC phenotype (32, 33). This pattern resembles the characteristic image of IH, where a gradual diminution of the vessel lumen occurs due to migration and proliferation of SMCs into the vascular intima (63). IH is the most frequent cause for vessel failure subsequent to aortocoronary vein grafting (64) and vascular interventions such as POBA and stent implantation (65), and the principal stimulus for IH appears to be endothelial injury. However, the exact mechanisms by which IH occurs are incompletely understood. Given the enhanced biomimetic features, our TEBV model is a promising approach towards an *in-vitro* system for the study of atherosclerosis and IH.

### TEBVs as a platform for intravascular device testing

O’Halloran-Cardinal et al. were the first to demonstrate that stent implantation as well as OCT imaging were feasible in TEBVs composed of an ePTFE scaffold with an intimal lining of human microvessel ECs (54). Some years later, the group was able to show accelerated regeneration of ECs over protein-modified stent strut surfaces with significantly increased tissue thickness compared to bare metal stents (55), supporting the use of TEBVs for evaluation of modified stent surfaces. Their model, however, lacked a medial layer, although some interspersed SMCs were noted below the endothelium, possibly because cells were isolated from liposuction fat, resulting in a mixed population of cells with mesenchymal and endothelial origin. Moreover, the compliance of the TEBV wall was limited due to the polymer scaffold employed to support the ECs. Therefore, this system has limited ability to fully mimic features of human blood vessels.

O’Cearbhaill et al. developed a more compliant TEBV with a backbone of silicone seeded with ECs which allowed for the incorporation of pressure, tensile hoop strain, and shear stress into the model (66). The group was able to demonstrate enhanced expression of genes indicating an inflammatory response in ECs at 24h after stent implantation, including E-Selectin, ICAM-1, and VCAM-1, along with an increased expression of the pro-apoptotic protein Bax and a decrease in the anti-apoptotic protein Bcl-2 and in Ki67. Although this model had improved ability to mimic hemodynamic forces, the absence of SMCs and ECM proteins within the vascular wall was a major limitation. Furthermore, the group used human umbilical vein ECs (HUVECs) instead of arterial ECs. However, ECs from different vascular beds have been shown to respond differently upon mechanical stimulation and possess distinct susceptibility towards atherosclerotic stimuli (67–69). Therefore, we chose ECs of arterial origin in our model.

### Inflammatory responses and endothelial healing following POBA and stent implantation in TEBVs

In line with findings from O’Cearbhaill et al. (66), we observed an inflammatory response in our model, as indicated by increasing levels of proinflammatory cytokines within the culture medium over the course of 12 days after stent implantation. Highest levels of IL-6, IL-8, and MCP-1 were observed in culture medium samples from BMS-treated TEBVs.

Mechanical injury, such as POBA and stent implantation, elicits an inflammatory response from the vessel wall, including the recruitment of monocytes and other leukocytes to areas of vascular injury, which is mediated by chemoattractant cytokines and interleukins (70–72). Cipollone et al. have demonstrated elevated levels of MCP-1 in patients with restenosis after POBA (73), and increased levels of IL-6 and IL-8 have been found after implantation of BMS and DES (74–76). However, comparisons of cytokine levels following BMS or DES-implantation in humans yielded conflicting results. The anti-proliferative drug-coating of DES (*i.e.*, everolimus in our study), possesses anti-inflammatory properties, as evidenced by their current use as immunomodulatory agents in the prevention and treatment of transplant rejection (77). Indeed, in a porcine model of stent implantation, Suzuki et al. found reduced expression of MCP-1 and IL-6 within the vessel wall after DES-implantation compared to the expression after BMS-implantation (78), and everolimus-eluting stents have been suggested to stabilize plaque inflammation *in vivo* (79). Other studies reported a local inflammatory response following first-generation DES-implantation (80, 81), and – compared with BMS-treated patients – systemic inflammatory responses were accentuated in patients undergoing DES-implantation for acute coronary syndrome (82). Nevertheless, while endothelial cells contribute to inflammatory reactions following vascular injury, the bulk of proinflammatory cytokines in humans is released by leukocytes. Thus, our model does not fully account for inflammation in its current form. Further studies including circulating lymphocytes are warranted to investigate inflammatory responses in TEBVs upon POBA and stent implantation.

In line with findings from clinical and histopathological studies (80, 83, 84), we observed delayed endothelial healing in TEBVs after DES-implantation, whereas more than 20% of stent struts were covered by neointima in BMS-treated TEBVs 12 days after the procedure. The antiproliferative drug-coating is intended to reduce neointimal formation by impeding SMC proliferation and migration, however, these drugs also impair the normal healing processes of the injured arterial wall (85–87).

Nevertheless, DES demonstrate a significant reduction in restenosis compared with BMS, and over the last decades, design and biocompatibility of DES have been evolving rapidly (88). Latest-generation DES are characterized by improved stent architecture with thinner strut backbones, increased radial strength and improved deliverability (89). They also have improved durable and biodegradable polymers – and in some cases no polymer – as well as new antiproliferative drug types and dosing. However, the issue of late in-stent restenosis, including neoatherosclerosis after DES implantation, is still warranted to overcome (90, 91). Further research should focus on strategies to minimize vascular injury and to improve biocompatibility of DES by material innovation, titration of drug-release and polymer degradation profile, and minimized strut thickness. Therefore, DES will continue to evolve, and TEBVs might potentially allow for the initial *in vitro* testing of novel stent devices.

### Limitations and Outlook

Our model currently recapitulates key features of intimal hyperplasia and provides a promising platform for future studies of atherosclerotic mechanisms. Atherosclerosis is a multifactorial disease, and while our model reproduces important aspects of atherogenesis-such as SMC migration and proliferation following endothelial injury - other critical components remain to be addressed. In particular, TEBVs were circulated with cell culture medium only, and thus were not exposed to the cellular and molecular constituents of human blood. Moreover, the hemodynamic properties of blood, includingturbulent flow and shear stress alterations, play a crucial role in atherogenesis. One key event in atherogenesis is the accumulation of lipids from the blood stream – mainly of LDL-C – within the vascular wall with subsequent leukocyte adhesion and their transmigration into the subendothelial space. Circulating blood cells are also thought to contribute to in-stent restenosis (92). Therefore, further investigations should aim to incorporate the cellular and molecular composition as well as the rheological properties of human blood to further advance the physiological relevance of the model.

Atherosclerosis, IH, and in-stent restenosis develop over years. However, maximum conditioning period was 21 days in our model. Therefore, the conditioning period should be extended, and further research should be undertaken to drastically accelerate lesion formation *in vitro*. This would allow for the *in vitro* testing of therapeutics that could directly target lesions in a controlled and measurable way.

We applied physiological flow and shear stress conditions to TEBVs. However, the VascuTrainer bioreactor system would potentially allow for the simulation of various hemodynamic conditions, including a disturbed flow, with oscillatory or recirculating flow conditions as found within areas in the vasculature that are prone to atherosclerosis and thrombus formation (93–95). Furthermore, we obtained cells from healthy subjects to generate TEBVs. In future experiments, the use of cells from patients with cardiovascular diseases would also be conceivable.

## Conclusion

Our three-dimensional TEBV model composed of SMCs and ECs recapitulates fundamental principles of vascular biology. An incomplete or absent endothelial layer resulted in increased proliferation of SMCs and collagen deposition. Thus, TEBVs represent a promising approach towards an *in vitro* model for the systematic investigation of IH and atherosclerotic pathways.

Moreover, POBA as well as the implantation of human-sized coronary stents was feasible with TEBVs in our model, as they are similar in size and wall thickness to human coronary arteries. Therefore, TEBVs have the potential to serve as a platform for high-throughput preclinical testing of cardiovascular devices.

## Abbreviation List

EC: endothelial cell
SMC: smooth muscle cell
ECM: extracellular matrix
POBA: plain old balloon angioplasty
IH: intimal hyperplasia
TEBV: tissue-engineered blood vessel
PVDF: polyvinylidene difluoride
BMS: bare metal stent
DES: drug-eluting stent
FB: fibroblast
OCT: optical coherence tomography
TBS: tris-buffered saline
αSMA: α-smooth muscle actin
IL-6: interleukin 6
IL-8: interleukin 8
MCP-1: monocyte chemoattractant factor 1

## a) Acknowledgements, Sources of Funding & Disclosures

Acknowledgements

We would like to acknowledge Mrs. Angela Freund for her invaluable technical assistance with embedding and slides production and Mr. Midhun Abraham Philip for assistance with OCT imaging.

## Highlights

- Current preclinical testing models do not adequately recapitulate the complexity of human arteries.
- Three-dimensional tubular constructs fabricated from fibrin-gel, supported by a textile mesh and seeded with ECs and SMCs enable extensive investigation of EC-SMC interactions under health and disease conditions.
- TEBVs can undergo vascular intervention including POBA and stent implantation and therefore would be useful for preclinical testing of coronary devices.

## References

1. Ross R. Atherosclerosis--an inflammatory disease. N Engl J Med. 1999;340(2):115–26.

2. Brunner H, Cockcroft JR, Deanfield J, et al. Endothelial function and dysfunction. Part II: Association with cardiovascular risk factors and diseases. A statement by the Working Group on Endothelins and Endothelial Factors of the European Society of Hypertension. J Hypertens. 2005;23(2):233–46.

3. Shi N, Mei X, Chen SY. Smooth Muscle Cells in Vascular Remodeling. Arterioscler Thromb Vasc Biol. 2019;39(12):e247–e52.

4. Lilly B. We have contact: endothelial cell-smooth muscle cell interactions. Physiology (Bethesda). 2014;29(4):234–41.

5. Li M, Qian M, Kyler K, Xu J. Endothelial-Vascular Smooth Muscle Cells Interactions in Atherosclerosis. Front Cardiovasc Med. 2018;5:151.

6. Méndez-Barbero N, Gutiérrez-Muñoz C, Blanco-Colio LM. Cellular Crosstalk between Endothelial and Smooth Muscle Cells in Vascular Wall Remodeling. Int J Mol Sci. 2021;22(14).

7. Sandoo A, van Zanten JJ, Metsios GS, Carroll D, Kitas GD. The endothelium and its role in regulating vascular tone. Open Cardiovasc Med J. 2010;4:302–12.

8. Fillinger MF, Sampson LN, Cronenwett JL, Powell RJ, Wagner RJ. Coculture of endothelial cells and smooth muscle cells in bilayer and conditioned media models. J Surg Res. 1997;67(2):169–78.

9. Powell RJ, Bhargava J, Basson MD, Sumpio BE. Coculture conditions alter endothelial modulation of TGF-beta 1 activation and smooth muscle growth morphology. Am J Physiol. 1998;274(2):H642–9.

10. Korff T, Kimmina S, Martiny-Baron G, Augustin HG. Blood vessel maturation in a 3-dimensional spheroidal coculture model: direct contact with smooth muscle cells regulates endothelial cell quiescence and abrogates VEGF responsiveness. Faseb j. 2001;15(2):447–57.

11. Pasternak RC, Baughman KL, Fallon JT, Block PC. Scanning electron microscopy after coronary transluminal angioplasty of normal canine coronary arteries. Am J Cardiol. 1980;45(3):591–8.

12. Block PC, Myler RK, Stertzer S, Fallon JT. Morphology after transluminal angioplasty in human beings. N Engl J Med. 1981;305(7):382–5.

13. Cornelissen A, Vogt FJ. The effects of stenting on coronary endothelium from a molecular biological view: Time for improvement? J Cell Mol Med. 2019;23(1):39–46.

14. Libby P. Molecular and cellular mechanisms of the thrombotic complications of atherosclerosis. J Lipid Res. 2009;50 Suppl(Suppl):S352–7.

15. Xu J, Shi GP. Vascular wall extracellular matrix proteins and vascular diseases. Biochim Biophys Acta. 2014;1842(11):2106–19.

16. Nagler A, Miao HQ, Aingorn H, Pines M, Genina O, Vlodavsky I. Inhibition of collagen synthesis, smooth muscle cell proliferation, and injury-induced intimal hyperplasia by halofuginone. Arterioscler Thromb Vasc Biol. 1997;17(1):194–202.

17. van Buul-Wortelboer MF, Brinkman HJ, Dingemans KP, de Groot PG, van Aken WG, van Mourik JA. Reconstitution of the vascular wall in vitro. A novel model to study interactions between endothelial and smooth muscle cells. Exp Cell Res. 1986;162(1):151–8.

18. Graham DJ, Alexander JJ, Miguel R. Aortic endothelial and smooth muscle cell co-culture: an in vitro model of the arterial wall. J Invest Surg. 1991;4(4):487–94.

19. Fillinger MF, O’Connor SE, Wagner RJ, Cronenwett JL. The effect of endothelial cell coculture on smooth muscle cell proliferation. J Vasc Surg. 1993;17(6):1058–67; discussion 67-8.

20. Wallace CS, Champion JC, Truskey GA. Adhesion and function of human endothelial cells co-cultured on smooth muscle cells. Ann Biomed Eng. 2007;35(3):375–86.

21. Ganesan MK, Finsterwalder R, Leb H, et al. Three-Dimensional Coculture Model to Analyze the Cross Talk Between Endothelial and Smooth Muscle Cells. Tissue Eng Part C Methods. 2017;23(1):38–49.

22. Chiu JJ, Chen LJ, Chen CN, Lee PL, Lee CI. A model for studying the effect of shear stress on interactions between vascular endothelial cells and smooth muscle cells. J Biomech. 2004;37(4):531–9.

23. Wolf F, Vogt F, Schmitz-Rode T, Jockenhoevel S, Mela P. Bioengineered vascular constructs as living models for in vitro cardiovascular research. Drug Discov Today. 2016;21(9):1446–55.

24. Barron V, Brougham C, Coghlan K, et al. The effect of physiological cyclic stretch on the cell morphology, cell orientation and protein expression of endothelial cells. J Mater Sci Mater Med. 2007;18(10):1973–81.

25. Fitzgerald KA, Malhotra M, Curtin CM, FJ OB, CM OD. Life in 3D is never flat: 3D models to optimise drug delivery. J Control Release. 2015;215:39–54.

26. Ryan AJ, Brougham CM, Garciarena CD, Kerrigan SW, O’Brien FJ. Towards 3D in vitro models for the study of cardiovascular tissues and disease. Drug Discov Today. 2016;21(9):1437–45.

27. Henderson AR, Choi H, Lee E. Blood and Lymphatic Vasculatures On-Chip Platforms and Their Applications for Organ-Specific In Vitro Modeling. Micromachines (Basel). 2020;11(2).

28. Wolf F, Rojas González DM, Steinseifer U, et al. VascuTrainer: A Mobile and Disposable Bioreactor System for the Conditioning of Tissue-Engineered Vascular Grafts. Ann Biomed Eng. 2018;46(4):616–26.

29. Mohapatra SR, Rama E, Melcher C, et al. From In Vitro to Perioperative Vascular Tissue Engineering: Shortening Production Time by Traceable Textile-Reinforcement. Tissue Eng Regen Med. 2022;19(6):1169–84.

30. Schindelin J, Arganda-Carreras I, Frise E, et al. Fiji: an open-source platform for biological-image analysis. Nature Methods. 2012;9(7):676–82.

31. Reddy GK, Enwemeka CS. A simplified method for the analysis of hydroxyproline in biological tissues. Clin Biochem. 1996;29(3):225–9.

32. Arnoldi R, Chaponnier C, Gabbiani G, Hinz B. Chapter 88 - Heterogeneity of Smooth Muscle. In: Hill JA, Olson EN, editors. Muscle. Boston/Waltham: Academic Press; 2012. p. 1183–95.

33. Hinz B. Myofibroblasts. Exp Eye Res. 2016;142:56–70.

34. Schmidt D, Asmis LM, Odermatt B, et al. Engineered living blood vessels: functional endothelia generated from human umbilical cord-derived progenitors. Ann Thorac Surg. 2006;82(4):1465–71; discussion 71.

35. Mallone A, Stenger C, Von Eckardstein A, Hoerstrup SP, Weber B. Biofabricating atherosclerotic plaques: In vitro engineering of a three-dimensional human fibroatheroma model. Biomaterials. 2018;150:49–59.

36. Kohlhaas J, Jäger MA, Lust L, De La Torre C, Hecker M, Korff T. Endothelial cells control vascular smooth muscle cell cholesterol levels by regulating 24-dehydrocholesterol reductase expression. Exp Cell Res. 2021;399(2):112446.

37. Wimmer RA, Leopoldi A, Aichinger M, Kerjaschki D, Penninger JM. Generation of blood vessel organoids from human pluripotent stem cells. Nat Protoc. 2019;14(11):3082–100.

38. Wimmer RA, Leopoldi A, Aichinger M, et al. Human blood vessel organoids as a model of diabetic vasculopathy. Nature. 2019;565(7740):505–10.

39. Dorweiler B, Torzewski M, Dahm M, et al. A novel in vitro model for the study of plaque development in atherosclerosis. Thromb Haemost. 2006;95(1):182–9.

40. Hsieh HJ, Li NQ, Frangos JA. Shear stress increases endothelial platelet-derived growth factor mRNA levels. Am J Physiol. 1991;260(2 Pt 2):H642–6.

41. Hsieh HJ, Li NQ, Frangos JA. Shear-induced platelet-derived growth factor gene expression in human endothelial cells is mediated by protein kinase C. J Cell Physiol. 1992;150(3):552–8.

42. Wang HQ, Huang LX, Qu MJ, et al. Shear stress protects against endothelial regulation of vascular smooth muscle cell migration in a coculture system. Endothelium. 2006;13(3):171–80.

43. Buga GM, Gold ME, Fukuto JM, Ignarro LJ. Shear stress-induced release of nitric oxide from endothelial cells grown on beads. Hypertension. 1991;17(2):187–93.

44. Garanich JS, Pahakis M, Tarbell JM. Shear stress inhibits smooth muscle cell migration via nitric oxide-mediated downregulation of matrix metalloproteinase-2 activity. Am J Physiol Heart Circ Physiol. 2005;288(5):H2244–52.

45. Rojas-González DM, Babendreyer A, Ludwig A, Mela P. Analysis of flow-induced transcriptional response and cell alignment of different sources of endothelial cells used in vascular tissue engineering. Scientific Reports. 2023;13(1):14384.

46. Korner PI, Angus JA. Vascular remodeling. Hypertension. 1997;29(4):1065–6.

47. van Varik BJ, Rennenberg RJ, Reutelingsperger CP, Kroon AA, de Leeuw PW, Schurgers LJ. Mechanisms of arterial remodeling: lessons from genetic diseases. Front Genet. 2012;3:290.

48. L’Heureux N, Pâquet S, Labbé R, Germain L, Auger FA. A completely biological tissue-engineered human blood vessel. Faseb j. 1998;12(1):47–56.

49. Kelm JM, Emmert MY, Zürcher A, et al. Functionality, growth and accelerated aging of tissue engineered living autologous vascular grafts. Biomaterials. 2012;33(33):8277–85.

50. Quint C, Arief M, Muto A, Dardik A, Niklason LE. Allogeneic human tissue-engineered blood vessel. J Vasc Surg. 2012;55(3):790–8.

51. Wystrychowski W, McAllister TN, Zagalski K, Dusserre N, Cierpka L, L’Heureux N. First human use of an allogeneic tissue-engineered vascular graft for hemodialysis access. J Vasc Surg. 2014;60(5):1353–7.

52. Peck M, Gebhart D, Dusserre N, McAllister TN, L’Heureux N. The evolution of vascular tissue engineering and current state of the art. Cells Tissues Organs. 2012;195(1-2):144–58.

53. Robert J, Weber B, Frese L, et al. A three-dimensional engineered artery model for in vitro atherosclerosis research. PLoS One. 2013;8(11):e79821.

54. Cardinal KO, Bonnema GT, Hofer H, Barton JK, Williams SK. Tissue-engineered vascular grafts as in vitro blood vessel mimics for the evaluation of endothelialization of intravascular devices. Tissue Eng. 2006;12(12):3431–8.

55. Cardinal KO, Williams SK. Assessment of the intimal response to a protein-modified stent in a tissue-engineered blood vessel mimic. Tissue Eng Part A. 2009;15(12):3869–76.

56. Punchard MA, O’Cearbhaill ED, Mackle JN, et al. Evaluation of human endothelial cells post stent deployment in a cardiovascular simulator in vitro. Ann Biomed Eng. 2009;37(7):1322–30.

57. L’Heureux N, Stoclet JC, Auger FA, Lagaud GJ, Germain L, Andriantsitohaina R. A human tissue-engineered vascular media: a new model for pharmacological studies of contractile responses. Faseb j. 2001;15(2):515–24.

58. Grassl ED, Oegema TR, Tranquillo RT. Fibrin as an alternative biopolymer to type-I collagen for the fabrication of a media equivalent. J Biomed Mater Res. 2002;60(4):607–12.

59. Long JL, Tranquillo RT. Elastic fiber production in cardiovascular tissue-equivalents. Matrix Biol. 2003;22(4):339–50.

60. Koch S, Flanagan TC, Sachweh JS, et al. Fibrin-polylactide-based tissue-engineered vascular graft in the arterial circulation. Biomaterials. 2010;31(17):4731–9.

61. Tschoeke B, Flanagan TC, Cornelissen A, et al. Development of a composite degradable/nondegradable tissue-engineered vascular graft. Artif Organs. 2008;32(10):800–9.

62. Fernández-Colino A, Wolf F, Rütten S, et al. Small Caliber Compliant Vascular Grafts Based on Elastin-Like Recombinamers for in situ Tissue Engineering. Front Bioeng Biotechnol. 2019;7:340.

63. Mills B, Robb T, Larson DF. Intimal hyperplasia: slow but deadly. Perfusion. 2012;27(6):520–8.

64. Wilbring M, Tugtekin SM, Zatschler B, et al. Even short-time storage in physiological saline solution impairs endothelial vascular function of saphenous vein grafts. Eur J Cardiothorac Surg. 2011;40(4):811–5.

65. Curcio A, Torella D, Indolfi C. Mechanisms of smooth muscle cell proliferation and endothelial regeneration after vascular injury and stenting: approach to therapy. Circ J. 2011;75(6):1287–96.

66. O’Cearbhaill ED, Punchard MA, Murphy M, Barry FP, McHugh PE, Barron V. Response of mesenchymal stem cells to the biomechanical environment of the endothelium on a flexible tubular silicone substrate. Biomaterials. 2008;29(11):1610–9.

67. Deng DX, Tsalenko A, Vailaya A, et al. Differences in vascular bed disease susceptibility reflect differences in gene expression response to atherogenic stimuli. Circ Res. 2006;98(2):200–8.

68. Sung HJ, Yee A, Eskin SG, McIntire LV. Cyclic strain and motion control produce opposite oxidative responses in two human endothelial cell types. Am J Physiol Cell Physiol. 2007;293(1):C87–94.

69. Rojas-González DM, Babendreyer A, Ludwig A, Mela P. Analysis of flow-induced transcriptional response and cell alignment of different sources of endothelial cells used in vascular tissue engineering. Sci Rep. 2023;13(1):14384.

70. Rollins BJ. Monocyte chemoattractant protein 1: a potential regulator of monocyte recruitment in inflammatory disease. Mol Med Today. 1996;2(5):198–204.

71. Welt FG, Rogers C. Inflammation and restenosis in the stent era. Arterioscler Thromb Vasc Biol. 2002;22(11):1769–76.

72. Webb LM, Ehrengruber MU, Clark-Lewis I, Baggiolini M, Rot A. Binding to heparan sulfate or heparin enhances neutrophil responses to interleukin 8. Proc Natl Acad Sci U S A. 1993;90(15):7158–62.

73. Cipollone F, Marini M, Fazia M, et al. Elevated circulating levels of monocyte chemoattractant protein-1 in patients with restenosis after coronary angioplasty. Arterioscler Thromb Vasc Biol. 2001;21(3):327–34.

74. Wang Z, Zhang T, Sun L, et al. Pioglitazone Attenuates Drug-Eluting Stent-Induced Proinflammatory State in Patients by Blocking Ubiquitination of PPAR. PPAR Res. 2016;2016:7407153.

75. Li X, Guo D, Zhou H, et al. Pro-inflammatory cytokines, oxidative stress and diffuse coronary reocclusions in elderly patients after coronary stenting. Cytokine. 2020;129:155028.

76. Wan M, Hu K, Lu Y, et al. Co-release of cytokines after drug-eluting stent implantation in acute myocardial infarction patients with PCI. Sci Rep. 2024;14(1):1236.

77. Tedesco-Silva H, Pascual J, Viklicky O, et al. Safety of Everolimus With Reduced Calcineurin Inhibitor Exposure in De Novo Kidney Transplants: An Analysis From the Randomized TRANSFORM Study. Transplantation. 2019;103(9):1953–63.

78. Suzuki T, Kopia G, Hayashi S, et al. Stent-based delivery of sirolimus reduces neointimal formation in a porcine coronary model. Circulation. 2001;104(10):1188–93.

79. Calfon Press MA, Mallas G, Rosenthal A, et al. Everolimus-eluting stents stabilize plaque inflammation in vivo: assessment by intravascular fluorescence molecular imaging. Eur Heart J Cardiovasc Imaging. 2017;18(5):510–8.

80. Joner M, Finn AV, Farb A, et al. Pathology of drug-eluting stents in humans: delayed healing and late thrombotic risk. J Am Coll Cardiol. 2006;48(1):193–202.

81. Wilson GJ, Nakazawa G, Schwartz RS, et al. Comparison of inflammatory response after implantation of sirolimus- and paclitaxel-eluting stents in porcine coronary arteries. Circulation. 2009;120(2):141–9, 1-2.

82. Liu J, Zhuo XZ, Liu W, et al. Drug-eluting stent, but not bare metal stent, accentuates the systematic inflammatory response in patients. Cardiology. 2014;128(3):259–65.

83. Nakazawa G, Ladich E, Finn AV, Virmani R. Pathophysiology of vascular healing and stent mediated arterial injury. EuroIntervention. 2008;4 Suppl C:C7–10.

84. Cornelissen A, Guo L, Fernandez R, et al. Endothelial Recovery in Bare Metal Stents and Drug-Eluting Stents on a Single-Cell Level. Arterioscler Thromb Vasc Biol. 2021;41(8):2277–92.

85. Marx SO, Jayaraman T, Go LO, Marks AR. Rapamycin-FKBP inhibits cell cycle regulators of proliferation in vascular smooth muscle cells. Circ Res. 1995;76(3):412–7.

86. Poon M, Marx SO, Gallo R, Badimon JJ, Taubman MB, Marks AR. Rapamycin inhibits vascular smooth muscle cell migration. J Clin Invest. 1996;98(10):2277–83.

87. Wiskirchen J, Schöber W, Schart N, et al. The effects of paclitaxel on the three phases of restenosis: smooth muscle cell proliferation, migration, and matrix formation: an in vitro study. Invest Radiol. 2004;39(9):565–71.

88. Torii S, Jinnouchi H, Sakamoto A, et al. Drug-eluting coronary stents: insights from preclinical and pathology studies. Nat Rev Cardiol. 2020;17(1):37–51.

89. Mori M, Sakamoto A, Sato Y, et al. Overcoming challenges in refining the current generation of coronary stents. Expert Rev Cardiovasc Ther. 2021;19(11):1013–28.

90. Madhavan MV, Kirtane AJ, Redfors B, et al. Stent-Related Adverse Events >1 Year After Percutaneous Coronary Intervention. J Am Coll Cardiol. 2020;75(6):590–604.

91. Sakamoto A, Sato Y, Kawakami R, et al. Risk prediction of in-stent restenosis among patients with coronary drug-eluting stents: current clinical approaches and challenges. Expert Rev Cardiovasc Ther. 2021;19(9):801–16.

92. Bayes-Genis A, Campbell JH, Carlson PJ, Holmes DR, Jr., Schwartz RS. Macrophages, myofibroblasts and neointimal hyperplasia after coronary artery injury and repair. Atherosclerosis. 2002;163(1):89–98.

93. Yoshida Y, Sue W, Okano M, Oyama T, Yamane T, Mitsumata M. The effects of augmented hemodynamic forces on the progression and topography of atherosclerotic plaques. Ann N Y Acad Sci. 1990;598:256–73.

94. Sumi T, Yamashita A, Matsuda S, et al. Disturbed blood flow induces erosive injury to smooth muscle cell-rich neointima and promotes thrombus formation in rabbit femoral arteries. J Thromb Haemost. 2010;8(6):1394–402.

95. Qiu J, Zheng Y, Hu J, et al. Biomechanical regulation of vascular smooth muscle cell functions: from in vitro to in vivo understanding. J R Soc Interface. 2014;11(90):20130852.

